# AnnotIEM: novel tool for microbiome species-level annotation of 16S gene based microbial sequencing

**DOI:** 10.1101/2024.10.31.621263

**Authors:** Madhumita Bhattacharyya, Luise Rauer, Matthias Reiger, Jamie Afghani, Claudia Huelpuesch, Claudia Traidl-Hoffmann, Avidan U. Neumann

**Affiliations:** Institute of Environmental Medicine and Integrative Health, Faculty of Medicine, University of Augsburg, Germany; Chair of Environmental Medicine, Technical University Munich, Augsburg, Germany; Institute of Environmental Medicine (IEM), Helmholtz Munich, Augsburg, Germany; Christine Kühne – Center for Allergy Research and Education (CK-Care), Davos, Switzerland; Institute for Food & Health (ZIEL), Technical University Munich, Freising, Germany

**Keywords:** microbial sequence annotation, taxonomic classification, species-level assignment next-generation sequencing, microbiome analysis, skin microbiome, gut microbiome

## Abstract

Identification of microbial species is essential for the generation of meaningful results from microbiome analysis. However, correct annotation of 16S microbial sequences remains a challenge, especially down to the species-level, because most available databases of 16S sequences are not well curated. AnnotIEM is a new tool for species-level annotation of sequences derived from 16S rRNA gene sequencing. AnnotIEM’s novel hit-selection algorithm combines top-hit and majority-hit alignment results from multiple databases. Benchmarking using in-vitro and in-silico mock community control datasets showed that AnnotIEM’s annotation precision on species-level (median=0.83) is significantly (p<0.0001) better than any currently available annotation tool. Benchmarking using a number of real-world environmental microbiome studies demonstrated that AnnotIEM gives higher fraction (58-68%) of species-level annotated sequences, as compared to any other currently available tool. In conclusion, AnnotIEM significantly improves bacterial annotation, which can greatly enhance the results of any microbiome study, and could be potentially used also for other organisms.

---

High throughput next-generation sequencing (NGS) has led to the development of culture and cloning free methods of microbiome analysis^1^. A common approach used for microbiome profiling includes extraction and amplification of 16S rRNA genes from a sample, followed by its sequencing. This is the most cost and labor effective method for the exploration of microbial communities from clinically relevant and/or environmental samples. To this date, many microbiome profiling protocols have been established for sequencing and identification of 16S full length or a selected variable region (e.g., V1-V3, V3-V4) of the 16S rRNA from bacteria. This small subunit ribosomal RNA gene has been sequenced for thousands of bacterial species and is accessible in publicly available databases, such as: Ribosomal Database Project (RDP)^2^, SILVA^3^, Greengenes^4^, EzBioCloud^5^, Genome Taxonomy Database (GTDB)^6^ and National Center for Biotechnology Information (NCBI) 16S rRNA Refseq^7^. Computational analysis of the sequence dataset begins with the merging and clustering of sequence reads into representative sequences, either by similarity (e.g., operational taxonomic units, OTUs^8^ or more recently and more accurately by denoising (e.g., amplicon sequencing variants, ASVs by DADA2^9^. These sequences are then annotated with their corresponding microbial taxonomy using the above publicly available databases. One of the crucial bottlenecks of this 16S rRNA based approach is the correct annotation of a relatively short stretches of 16S rRNA, which becomes especially difficult due to the fact that the available 16S sequence databases are mostly not well curated and contain many mistakes.

The most problematic feature of 16S sequence taxonomy identification is annotation on the microbial species level. However, knowing the species of the various sequences in a given microbial community is of outmost importance both clinically and scientifically. For example, high relative abundance of *Staphylococcus aureus* in lesional skin is an indicator of severe atopic dermatitis (AD) disease, whereas other *Staphylococci* species such as *S. epidermidis* or *S. hominis* may be beneficial on the skin of AD patients^10-12^. Another important example is that while *Helicobacter pylori* causes peptic ulcers and gastric cancers, it nevertheless reduces the risk of inflammatory bowel disease (IBD), while other enterohepatic *Helicobacter* species increase the risk of IBD^13^. Also, dental caries was shown to be caused by *Streptococcus mutans*^14^, while colonization by *Streptococcus oligofermentans* gives rise to a reduced incidence of caries^15^. However, currently available annotation tools either do not give species level annotation at all or only give it for a small fraction of the sequences found.

Currently, the most commonly used tools for merging and clustering sequences are QIIME2^16^, DADA2^9^, IMNGS^17^, and Mothur^18^, which all have integrated annotation tools (Table 1). The IMNGS server uses an integrated RDP classifier with the SILVA database and provides only genus level annotation. DADA2 provides additional tools integrated in the package that assigns taxonomy to sequences by aligning them to Silva or Greengenes or the combination of RDP with NCBI sequence databases. Mothur includes a number of integrated taxonomy classifiers including the Basic Local Alignment Search Tool (BLAST) suite or K-mer based Wang algorithm, which can align sequences against RDP^2^, SILVA^3^ or Greengenes databases^4^. The Q2Classifier plugin^19^, the integrated classifier program in QIIME2^16^, includes many different classifier options such as nucleotide BLAST (BLASTn), RDP classifier, UCLUST classifier, SortmeRNA, VSEARCH etc. It is possible to align OTUs and ASVs against SILVA, Greengenes or any database formatted for QIIME2 using any of these classifiers. Another program, CLC workbench, which is distributed with a commercial license by Qiagen, uses in-house multiple sequence alignments for assigning taxonomy. However, the fraction of sequences that are successfully annotated on the genus level is often quite low for each of these programs, and even lower for species level annotation if possible at all.

**Table 1:** Properties of widely used annotation tools used for benchmarking of the AnnotIEM tool.

|  | DADA2-taxonomy | QIIME2 | Silva ACT | RDP SeqMatch | EzBiocloud | NCBI | AnnotIEM |
| --- | --- | --- | --- | --- | --- | --- | --- |
| **Database used** | Silva or Greengenes or RDP+NCBI | Green-genes or Silva or others | Silva | RDP | EzBiocloud | NCBI 16s rRNA database | Silva + RDP + GTDB + NCBI 16s rRNA DB |
| **Alignment algorithm** | RDP Classifier | BLAST VSEARCH SK-learn | K-MER based inhouse program | K-MER based inhouse program | BLASTn followed by Mayers and Millers program | BLASTn | BLASTn followed by AnnotIEM algorithm |
| **Server / Standalone** | Standalone | Standalone | Server | Server | Server | Server | Standalone |
| **License type** | Open source | Open source | Free | Free | Free/ Commercial* | Free | Open source |
| **Programing skill needed** | Moderate | High | Low | Moderate | Low | Moderate | Low |
| **Batch run capacity** | Yes | Yes | Yes (<1000) | Yes (<800) | No* | Yes | Yes |

\* Free for running individual sequences; Batch run requires a commercial license

Most of the above-mentioned programs require a moderate to high level of computational skill and often require Linux operating system. Another option for annotation is to use online servers such as Silva ACT^20^, RDP Seqmatch^21^, NCBI^22^, or EzBioCloud^5^ (Table 1). Silva ACT^20^ and RDP Seqmatch^21^ use an in-house K-mer based alignment algorithm with their corresponding databases. RDP Seqmatch typically gives species level annotation for about 20% of sequences, while Silva ACT only provides genus level annotation. NCBI uses BLASTn^22^ for alignment with the NCBI highly curated 16S rRNA database for providing species level annotation. EzBioCloud developed an algorithm using BLASTn^22^ followed by a global alignment with Mayers and Miller algorithm in combination with a propriety highly curated database, and gives better annotation results. However, while this platform is freely available for academic users, annotating more than one sequence at a time is possible only with a commercial license.

The quality of annotation by any of these tools is strongly dependent on the databases used, where either SILVA, RDP, Greengenes or NCBI is implemented. The first two databases are semi curated and the last two are more carefully curated. All these databases are updated semi-regularly. However, the usage of only one database often does not provide not provide an adequate coverage of species level annotation for all sequences. Furthermore, because most above databases contain either some erroneous entries or entries with meaningless information (e.g. “bacteria from arctic water”), it often happens that the top alignment hit gives an entry with missing or erroneous information. Thus, here we developed and benchmarked a new tool - AnnotIEM, which aligns microbial sequences against a number of different databases and provides species level annotation using a novel algorithm that utilizes both the top hit and the majority hit from the multiple databases used.

## Results

### Combination of top hit and majority hit from several databases

The core of the AnnotIEM algorithm (Figure 1A and Supplementary Figure S1) is based on the combination of the alignment results from several microbial sequence databases (by default, SILVA, RDP, GTDB and NCBI), and checking both the top hit (hit with best alignment scores) and the majority hit (species, or genus, with highest number of hits) together from all databases. BLASTn alignment results for the given 16S rRNA gene sequence (ASV or OTU) from each of the databases are parsed and filtered separately in order to identify the top ten hits from each database. Then a pool of all these hits are analyzed together by a novel algorithm that selects the overall top hit as well as the majority hit from all databases and uses alignment scores and a number of parameters (see details in methods) to identify the best possible species or genus level annotation.

**Figure 1:**
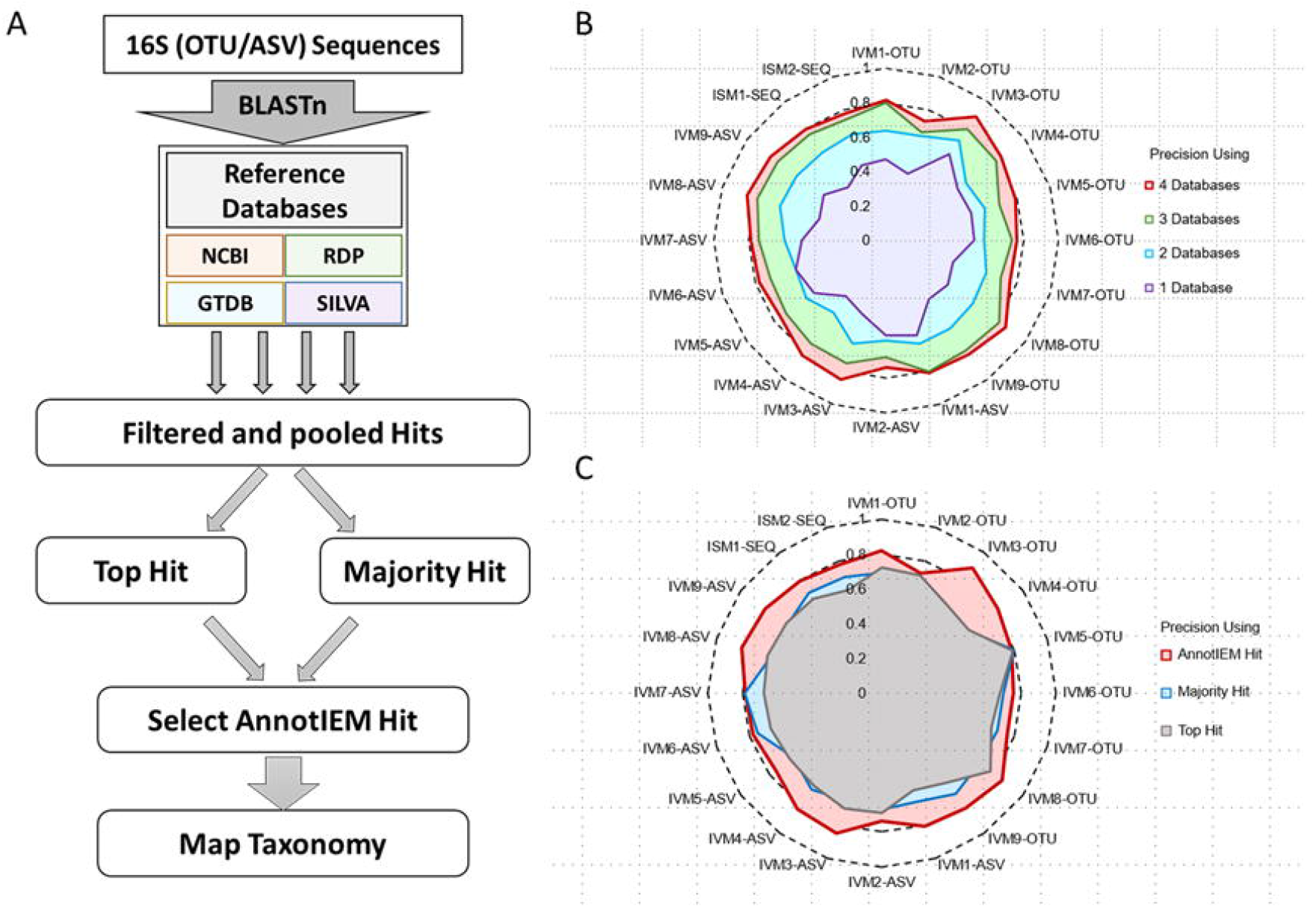
Algorithm overview and precision of species-level annotation by AnnotIEM using top-hit and majority-hit from four databases. A) Flowchart of the AnnotIEM algorithm explaining how a number of databases is used to produce top-hit and majority-hit annotation, followed by the selection of the final annotation (see Supplementary Figure S1 describing the detailed algorithm for selection of top-versus majority-hit). B) Precision of the AnnotIEM annotation on species-level in 9 in-vitro mock datasets (IVM), tested both as ASVs and OTUs, and in 2 in-silico mock (ISM) datasets (Table 2 and Supplementary Table S1). The radar chart shows the precision of annotation when using 1, 2, 3 or 4 databases, where the best results are for 4 databases, as used in the AnnotIEM tool (the full results for any combination of databases are shown in Supplementary Figure S2). C) Precision of the AnnotIEM annotation on species-level in the above 20 datasets when using 4 databases for only top-hit, only majority-hit, or the combination of top-hit and majority-hit with the AnnotIEM algorithm, which gives the best results.

In order to test AnnotIEM’s performance, we used sequence (both ASVs and OTUs) datasets from nine in-vitro mock (IVM, total of 189 sequences) communities along with sequences from two in-silico mock (ISM, each with 259 species) communities (details in Table 2 and Supplementary Table S1). The precision of species-level annotation using AnnotIEM with four databases was higher than 0.8 in 50% of the datasets and higher than 0.7 in all datasets (Figure 1 and supplementary Figure S2). Most of the errors, mock sequences that were not annotated correctly by AnnotIEM on species-level as compared to the original species, resulted from cases were there is a pair of species from the same genus with high (>99%) identity in the 16S region that was sequenced (e.g., *Clostridium beijerinckii* and *Clostridium acidotolerence* have 99% identity in their 16S V1-V3).

**Table 2:**
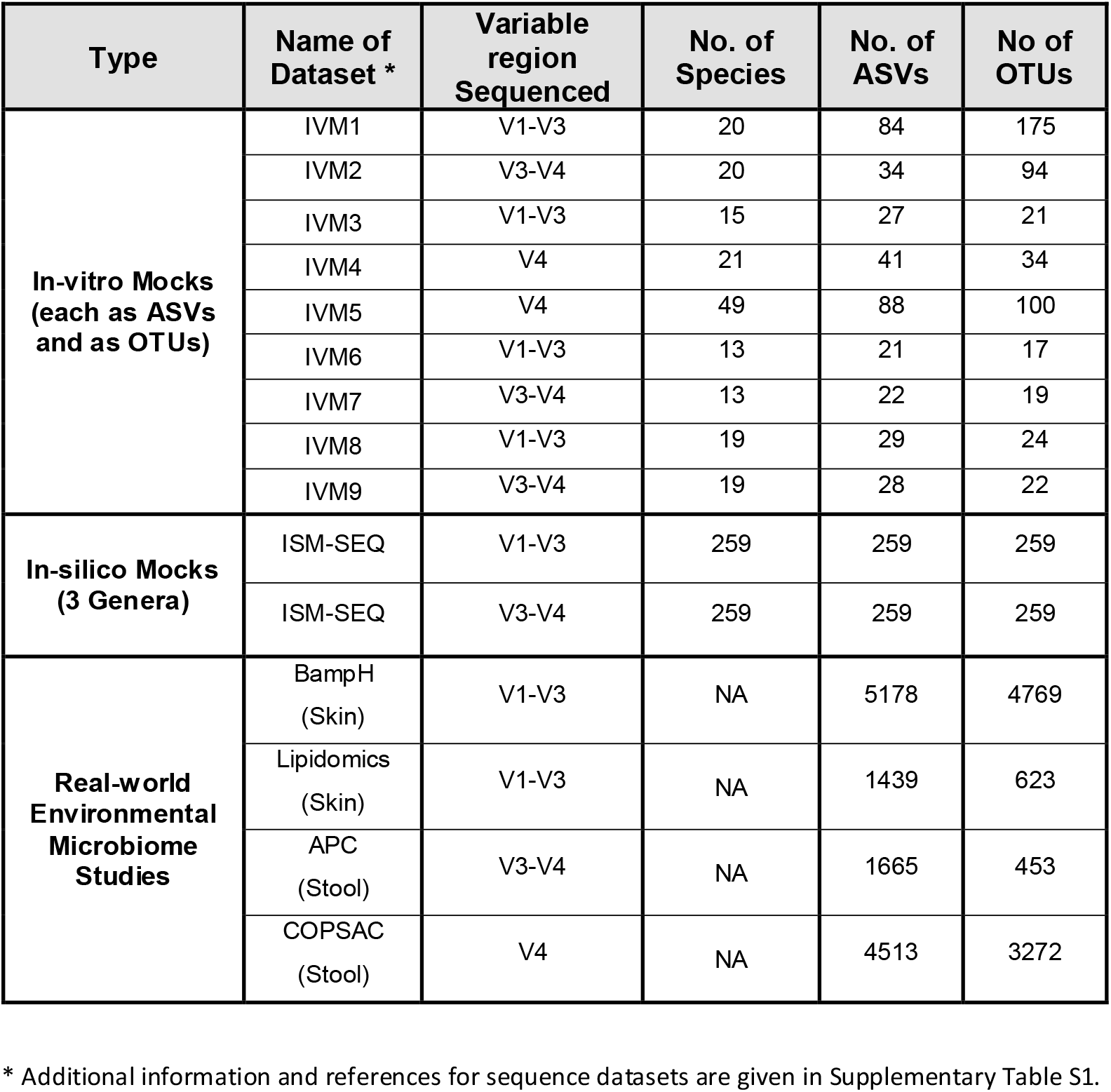
Sequence datasets used for benchmarking.

| Type | Name of Dataset * | Variable region Sequenced | No. of Species | No. of ASVs | No of OTUs |
| --- | --- | --- | --- | --- | --- |
| **In-vitro Mocks**<br>**(each as ASVs**<br>**and as OTUs)** | IVM1 | V1-V3 | 20 | 84 | 175 |
|  | IVM2 | V3-V4 | 20 | 34 | 94 |
|  | IVM3 | V1-V3 | 15 | 27 | 21 |
|  | IVM4 | V4 | 21 | 41 | 34 |
|  | IVM5 | V4 | 49 | 88 | 100 |
|  | IVM6 | V1-V3 | 13 | 21 | 17 |
|  | IVM7 | V3-V4 | 13 | 22 | 19 |
|  | IVM8 | V1-V3 | 19 | 29 | 24 |
|  | IVM9 | V3-V4 | 19 | 28 | 22 |
| **In-silico Mocks**<br>**(3 Genera)** | ISM-SEQ | V1-V3 | 259 | 259 | 259 |
|  | ISM-SEQ | V3-V4 | 259 | 259 | 259 |
| **Real-world**<br>**Environmental**<br>**Microbiome**<br>**Studies** | BampH<br>(Skin) | V1-V3 | NA | 5178 | 4769 |
|  | Lipidomics<br>(Skin) | V1-V3 | NA | 1439 | 623 |
|  | APC<br>(Stool) | V3-V4 | NA | 1665 | 453 |
|  | COPSAC<br>(Stool) | V4 | NA | 4513 | 3272 |

\* Additional information and references for sequence datasets are given in Supplementary Table S1.

Furthermore, for all data sets, using AnnotIEM with four databases resulted in significantly (p<0.001) much higher Precision (median 0.83) than using only any one of the databases (median 0.36), or any combination of two (median 0.54) or three (median 0.70) databases (Figure 1B and Supplementary Figure S2). Furthermore, using AnnotIEM with the combination of both top hit and majority hit gave higher annotation precision than using only the top hit or only the majority hit in 90% of the datasets (Figure 1C).

### Benchmarking AnnotIEM against other tools in control mock datasets

Next, we benchmarked AnnotIEM against other currently available annotation tools (Table 1) using the 11 control mock datasets (Table 2 and Supplementary Table S1). AnnotIEM demonstrated significantly more accurate annotation in comparison to all other tools, both on the species level annotation, for those tools that allow at all species level annotation, and on the genus level annotation (Figure 2 for ASVs and Supplementary Figure S3 for OTUs).

**Figure 2:**
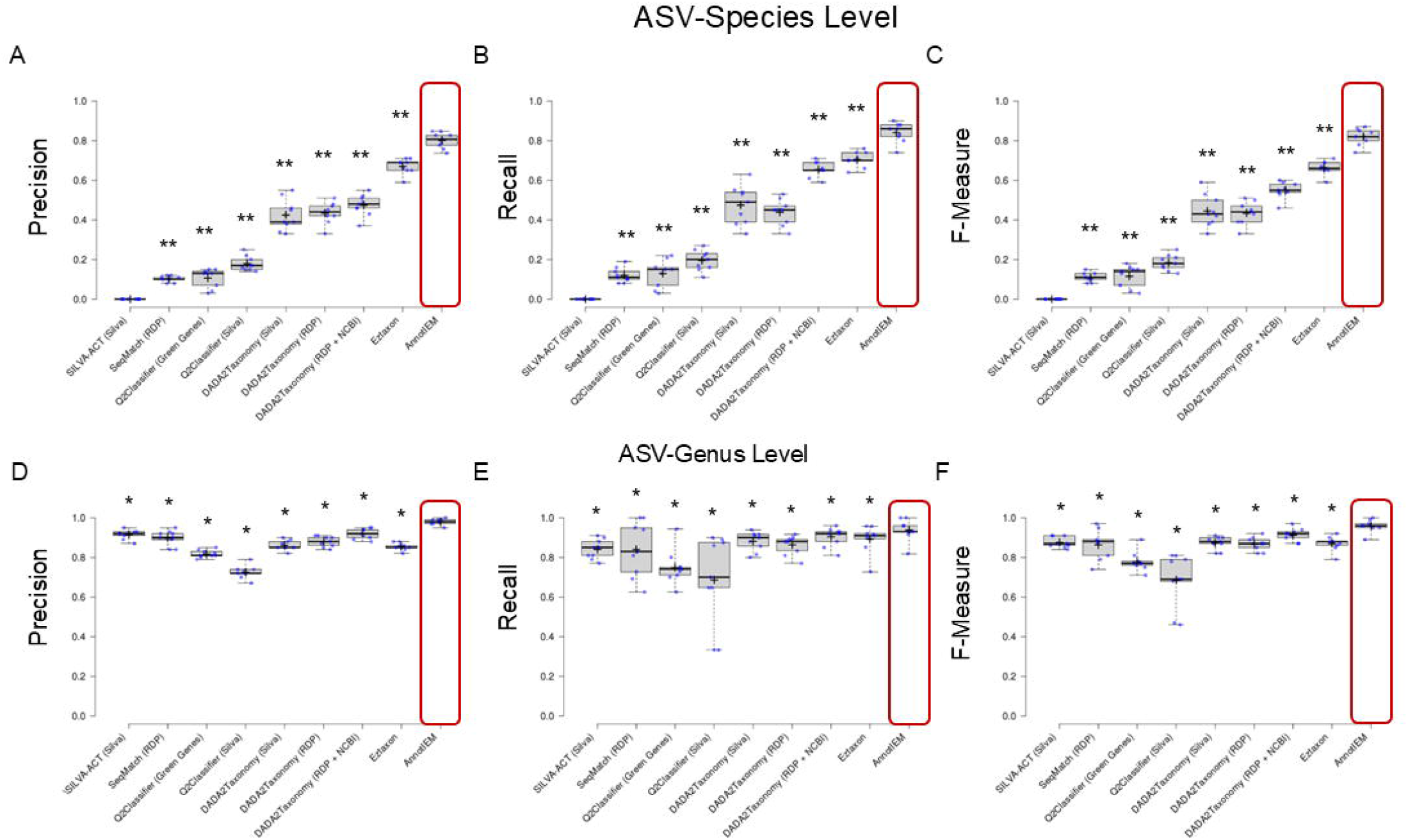
Benchmarking the annotation precision of AnnotIEM against other commonly used annotation tools using ASVs from mock communities. The performance of AnnotIEM is benchmarked against the most widely used annotation tools, using ASVs derived from 9 in-vitro mock (IVM) datasets (Table 2 and Supplementary Table S1). Scoring of the annotation on species-level (A-C) and on genus-level (D-F) is done by Precision (A and D, accordingly), Recall (B and E, accordingly) and F-Measure (C and F, accordingly) scores. The distribution of scores is shown as a box-whisker and scatter plot for each tool. Each box represents the 25th to 75th percentile of the distribution and whiskers extend to the 5th to 95th percentile. The mean is marked by a cross sign, and the median is marked by a horizontal line. * depicts p-value of p<0.05. ** depicts p-value of p<0.01. Commonly used tools used for benchmarking were: SILVA-ACT, SeqMatch RDP, Q2Classifier from QIIME2 using either the Greengenes or the SILVA database, DADA2taxonomy using either SILVA, RDP, or RDP+NCBI databases, and EzBioCloud.

On species level (Figure 2A-C), the Precision achieved by AnnotIEM for all mock communities (median=0.83) was significantly higher (p<0.0001) than all other tools (median=0.21 to 0.71). A similar result was observed for the Recall and F-measure scores. The second-best results were given by the EzBioCloud online tool, which notably allows only annotation of one sequence at a time when using the free version.

Annotation on the genus level by existing tools is considerably better than annotation on species level (Figure 2D-F). Nevertheless, the precision of genus level annotation by AnnotIEM was still significantly (p<0.05) higher (Median=0.96) as compared to the precision by the other tools (Median=0.52 to 0.88).

### Benchmarking AnnotIEM against other tools in environmental microbiome studies

Furthermore, we benchmarked AnnotIEM against other currently available annotation tools (Table 1) in four environmental stool and skin microbiome datasets (Table 2). The fraction (coverage) of 16S rRNA sequences annotated to species or genus level was used to compare the performance of the tools (Figure 3 for ASVs and supplementary Figure S4 for OTUs). In all four datasets AnnotIEM species level annotation was possible for at least 58% of sequences (58% to 68%), whereas only 42% or less (0% to 42%) of the sequences were annotated at species level by any of the other tools, if given at all (Figure 3A-D). AnnotIEM resulted also in better coverage on the fraction of sequences annotated on the genus level, although the performance of some of the other tools on genus level was not much lower. Note that we did not benchmark the EzBioCloud tool for the environmental datasets because of its limitation of sequencing only one sequence at a time in its free version.

**Figure 3:**
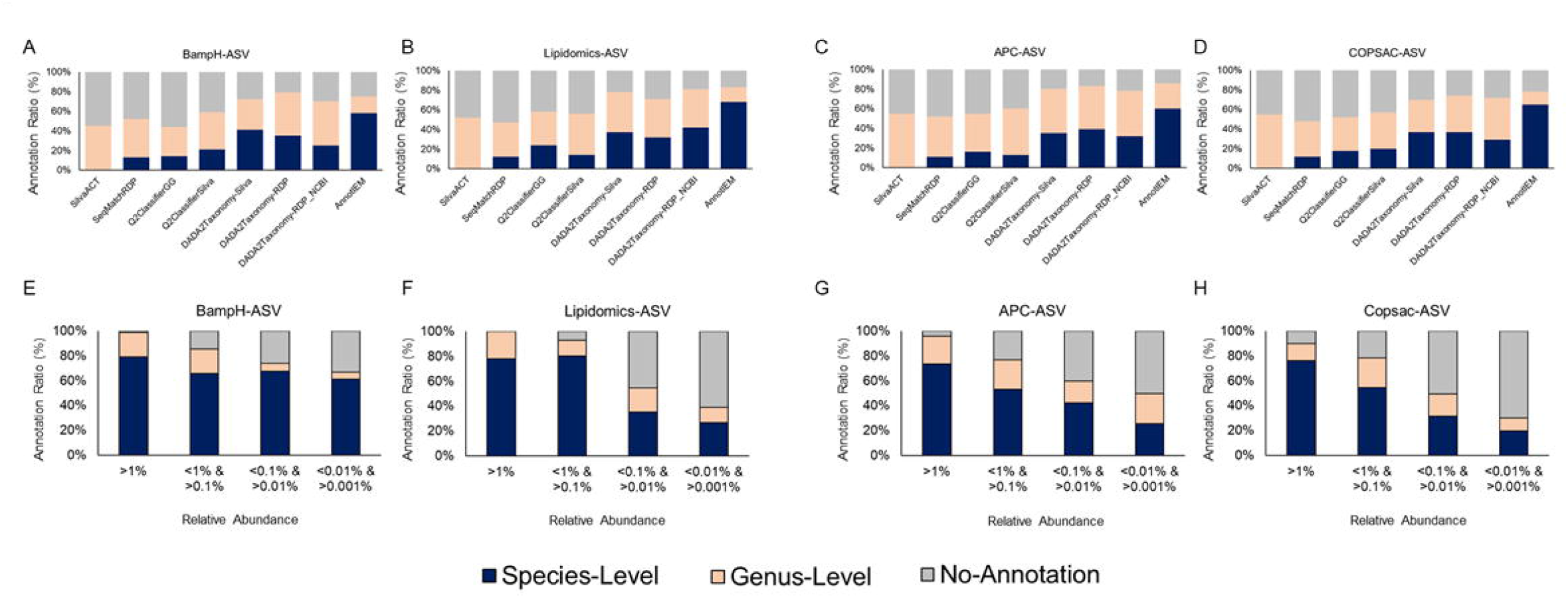
Benchmarking the species-level annotation coverage of AnnotIEM against other commonly used annotation tools using ASVs from real-world microbiome studies. The species-level, and genus-level, annotation coverage (fraction of sequences with species-level, and genus-level, annotation) of AnnotIEM is benchmarked against the most widely used annotation tools, using ASVs derived from 4 real-world large skin or gut microbiome case studies: BampH (A), Lipidomics (B), APC (C), and COPSAC (D), see additional information in Table 2 and Supplementary Table S1. For each of the studies, the species-level and genus-level annotation coverage by AnnotIEM is also depicted as function of the ASVs relative abundance (E-H). Commonly used tools used for benchmarking were: SILVA-ACT, SeqMatch RDP, Q2Classifier from QIIME2 using either the Greengenes or the SILVA database, and DADA2taxonomy using either SILVA, RDP, or RDP+NCBI databases. EzBioCloud tool was not benchmarked here because of its limitation of sequencing only one sequence at a time in its free version.

Interestingly, we observed that the annotation coverage by AnnotIEM increases as a function of the relative abundance of the sequences in the four environmental microbiome datasets (Figure 3E-H for ASVs and supplementary Figure S4E-H for OTUs). In all four datasets, ASVs with a relative abundance larger than 1% have a high coverage (73% to 79%) on the species level, and even higher up to genus level (86% to 98%), while ASVs with lower and lower relative abundance show lower and lower coverage on species and genus level.

### AnnotIEM added-value as a user-friendly tool

Last but not least, it is important to mention that AnnotIEM is a relatively user-friendly tool, which is simple to run with an easy to understand output. Also, the installation of AnnotIEM is straightforward (either on a Linux machine or on a virtual Linux machine running in other environment), since we give preformatted databases with the tool. Advanced users can change the databases used, either to use more updated versions of the databases used by default or use other databases, with instructions given in the README file of AnnotIEM.

Another important aspect of AnnotIEM as a tool is that it generates as result files both a simple to read summary of the final species (or genus) level annotation selected by AnnotIEM per sequence, and a more detailed result file with a description of the top hit and majority hit at both genus and species level from all four databases and the relevant annotation parameters per sequence. The latter detailed file can be used by advanced users to inspect more carefully sequences of special interest.

## Discussion

We introduce here a new tool – AnnotIEM – for precise species level annotation of microbial 16S rRNA gene sequences, which, by benchmarking against several of the currently available annotation tools, shows significantly better annotation, both in precision and in coverage. AnnotIEM achieves better annotation by using a novel algorithm that selects the best annotation based on the combination of top hit and majority hit from several databases. This is most probably due to inherent errors and missing information in large-scale semi curated biological sequence databases, where a large proportion of sequences lacks a full and/or correct annotation. Comparing the top hit with majority hit from several databases adds a level of confidence to this heuristic search results. Thus, this method minimizes, albeit does not completely negates, the chance of including a missed/incomplete/wrong annotation available from certain databases for certain sequences.

AnnotIEM performs better than all the other tools tested here, when compared on the species level annotation. Only the EzBioCloud tool performed almost as well, probably because of its use of a propriety curated database, but notably the free version of EzBioCloud allows annotating only one sequence at a time and thus is not practical for large datasets. It is worth to mention that also in genus level AnnotIEM gave better annotation than the other tools tested here, but there the improvement was less striking.

While the use of AnnotIEM together with algorithms for denoising of sequence data (e.g., DADA2 giving ASVs^9^) considerably increases the accuracy and coverage of annotation on microbial species level, still the identification of species with closely similar sequences (e.g., *Staphylococcus aureus* and *Staphylococcus simae*^23^ is still a challenge. AnnotIEM successfully annotated closely related species in all the mock communities (Table 2), with only a small number of mistakes due to the fact that some species have a 16S variable region identity larger than 99%. Moreover, the quality of annotation depends also on the 16S region (e.g., V1-V3, V3-V4 or V9) selected for sequencing, since different species are more differentiable on different 16S regions^24^. An extensive analysis is still needed to evaluate how sequencing different variable regions of the 16S gene may influence the identification of closely similar species. Supplying the users with a table of specific pairs of species annotations that may be misleading due to their almost identical sequence in different 16S variable regions would be highly beneficial. Nevertheless, AnnotIEM gives as an output both a shorthand version of the annotation results and a more detailed annotation results output that includes all parameters used for the annotation, which can be used by advanced users to evaluate how good the annotation of specific sequences of interest is.

As a tool AnnotIEM is user-friendly since the user needs to only run a one-line command from a Linux terminal, once the BLAST suite and sequence databases are installed. While we have tested AnnotIEM with four databases (RDP^2^, Silva^3^, NCBI 16S rRNA Refseq^7^ and GTDB^6^), the number and identity of the databases used can be modified by advanced users. Concluding from our tests with 1-4 databases, the use of even more, and better curated, databases, will very likely further improve the annotation results.

Currently AnnotIEM is optimized and tested only for bacterial 16S rRNA gene sequences; however, the algorithm can be implemented for any organism by altering the reference databases, for example for fungal 18S rRNA gene sequences or even for the newly developed databases and method for RNA virus annotation using the RdRP sequences^25^. In general, the novel annotation method introduced by AnnotIEM, combining the top hit and majority hit from several databases, should improve the annotation of any organism sequences in more detailed taxonomic level.

In conclusion, AnnotIEM is a valuable tool for microbiome analysis that by increasing the precision and coverage of species-level annotation will allow microbiome studies a better scientific and clinical interpretation.

## Online Methods

### Algorithm

AnnotIEM algorithm essentially runs in two parts. In the first part the query sequence sets are aligned by BLASTn against each of the reference databases separately. Here we used the following databases: RDP^2^, SILVA^3^, NCBI 16S rRNA Refseq^7^, and GTDB^6^. For each sequence for each database the alignment parameters: evalue ≤10^-5^, qCov ≥95%, and identity ≥95%, are used for filtering the hits. Results from each BLASTn run are parsed separately and the top 10 hits (based on identity) from each database are combined together.

In the second part of the algorithm, the combined hits are ranked according to identity value and the best match (highest identity) with species-level annotation is defined as the total species-level top-hit. All the other combined hits from all databases with species-level annotation are grouped to identify the species-level majority-hit, being the species with the maximum number of hits among the combined hits.

Then, a novel algorithm is used for selecting the best possible species-level annotation between the top-hit and majority-hit. For this algorithm, three different conditions, namely: difference in identity<dI, difference in rank <dR, and frequency of the majority-hit >F_m_, are used together (Supplementary Figure S1) to select as final species-level annotation either the top-hit or the majority-hit.

If species-level annotation is not available among the top 10 hits from all databases then the above algorithm is performed for genus-level annotation. If also no genus-level annotation is available, then AnnotIEM will report the result for this sequence as “No-Annotation”.

Finally, the full taxonomic classification hierarchy for the selected annotation is mapped. If no taxonomic classification can be retrieved, for example in case that the selected top-hit has an entry such as “Arctic water bacteria”, then this sequence will be marked as “Problematic”.

Note that AnnotIEM does not attempt to resolve whether the entries of the top 10 hits per database are valid microbial taxa or not. Also, AnnotIEM does not resolve multiple names used for certain microbial taxa (e.g., *Cutibacterium acnes* and *Propionibacterium acnes*), thus AnnotIEM will consider and report such cases as two different taxa.

### Tool availability

The tool and four databases needed for the default use of AnnotIEM, as well as a README instructions file for installation and running the tool and example datasets (used here for benchmarking), are available for download in https://github.com/Madhu84life/AnnotIEM.git

### Datasets used for benchmarking

The performance of AnnotIEM in comparison to other tools was benchmarked using three different types of datasets (Table 2 and Supplementary Table S1). A) In-vitro Mocks (IVM): Representative ASV (and OTU) sequences (total 189 sequences) from nine biological mock communities, among which four were kindly provided by our collaborators from ZIEL^26^. One mock community was cultured and extracted at the Institute of Environmental Medicine. The remaining four were retrieved from publicly available databases^27-29^. B) In-sillico Mocks (ISM): A synthetic mock community was constructed for 2 different regions (V1-V3 and V3-V4) of the 16S rRNA gene (see below), each with 259 species from three different genera: *Staphylococcus, Streptococcus* and *Clostridium*. C) Four environmental case studies of stool and skin microbiome from different projects^30,31^are also used for benchmarking analysis. Detailed information about all these datasets is provided in Supplementary Table S1.

### In-vitro mock communities

A total of nine in-vitro mock communities have been used in this study. Raw fastq sequence files were demultiplexed and processed by standardized pipelines to obtain ASVs (using DADA2) and OTUs (using Vsearch through QIIME2). The dataset also contains one in-house mock community created by *in-vitro* culture mix sequenced at ZIEL and processed to obtain OTUs (using IMNGS) and ASVs (using DADA2) [NCBI accession: https://www.ncbi.nlm.nih.gov/bioproject/PRJNA674596 ].

### In-silico mock communities

All the available full-length sequences from three bacterial genera (*Staphylococcus, Streptococcus* and *Clostridium*) were collected from the NCBI sequence database ^7^An in-house computational program was used to extract the V1-V3 (primer 27F-534R) and V3-V4 (337F-805R) regions. A total of 260 sequences were obtained for each of the two synthetic mock communities.

### Environmental microbiome studies

Four different case studies (both published and unpublished) have been used for benchmarking AnnotIEM:

1. Lipidomics: Skin microbiome sampled from atopic dermatitis (n=16) and acne patients (n=21) and matching healthy controls. Samples were sequenced at ZIEL and representative sequence sets were generated by IMNGS (OTUs) and DADA2 (ASVs). [ENA accession: https://www.ebi.ac.uk/ena/browser/view/PRJEB66070 ].
2. BampH: Skin microbiome sampled from atopic dermatitis patients (n=6) and healthy controls (n=6). Samples were sequenced at IMGM and representative sequence sets were generated by CLC workbench (OTUs) and DADA2 (ASVs). [ENA accession: https://www.ebi.ac.uk/ena/browser/view/PRJEB37663 ].
3. Atopy in Preterm Children (APC): This is a longitudinal study of stool microbiome in 53 children from 0 to 3 years. Samples were sequenced at ZIEL and representative sequence sets were generated by IMNGS (OTUs) and DADA2 (ASVs).
4. COPSAC: Stool microbiome samples from babies over three time points at: 1 week, 1 month and 1 year. The samples were collected, prepared and extracted as par the protocol of the COPSAC 2010 cohort. [NCBI accession: https://www.ncbi.nlm.nih.gov/bioproject/PRJNA417357 ].

### Benchmarking

Sequences (ASVs and OTUs) from all the above-mentioned datasets were annotated by AnnotIEM. For benchmarking against existing tools (both stand alone and online servers), each of the dataset was also annotated by online servers such as SINA tools from Silva-ACT^20^, Seqmatch tool from RDP^21^, EzBioCloud^5^, and NCBI-BLASTn^22^. Standalone tools such as the Q2Classifier^19^ plugin from QIIME2 using either SILVA or Greengenes as databases and the DADA2-assignTaxonomy function using either SILVA, RDP or RDP and NCBI combined as databases were used to annotate the above-mentioned datasets.

### Scores for annotation performance

For each mock dataset, the species used in each mock dataset were considered as the expected list of annotations. Each dataset was annotated individually with multiple tools. For each tool, the annotated list of bacteria was considered separately as observed annotation for the particular tool.

The annotation results were classified into: true positive (TP) annotations present in both expected and observed sets, false positive (FP) annotations that were observed but not expected, and false negative (FN) annotations that were expected but not observed. Note that true negative (TN) annotations are intrinsically not possible when using a known mock dataset. Using these three annotation categories, the following scores were calculated for each of the biological and synthetic mock communities:

Precision = TP / (TP + FP), Recall= TP / (TP + FN),

F-measures = 2 × precision × recall / (precision + recall).

For each of the environmental microbiome case datasets, the fraction of annotated sequences on species-level (or on genus-level) out of all sequences for each dataset were used to estimate the annotation coverage on species-level (or on genus-level). Of course, since the expected sequences are not known in the real-world microbiome studies, the annotation accuracy cannot be estimated for these datasets.

## Acknowledgments

We are grateful for Klaus Neuhaus and Sandra Reitmeier from ZIEL (Technical University of Munich) for providing mock datasets, as well as Mathis Hjelmsø, Jakob Stockholm and Casper Poulsen (University of Copenhagen) for providing the COPSAC microbiome dataset. We thank all above researchers and Christian Müller (Helmholtz Munich) for fruitful discussions.

## Author Contributions

Conceptualization: MB, MR and AUN; Methodology: MB and AUN with contributions from LR and CH; Software coding: MB; Data curation: MB; Data analysis: MB with contributions from LR and AUN; Beta testing: LR, JA and CH; Funding acquisition: MR and CTH; Visualization: MB and AUN with contribution from LR, MR, JA and CH; Writing—original draft: MB and AUN; Writing—review and editing: LR, MR, JA, CH and CTH. All authors have read and agreed to the published version of the manuscript.

## Supplementary Material

**Supplementary Table S1:**
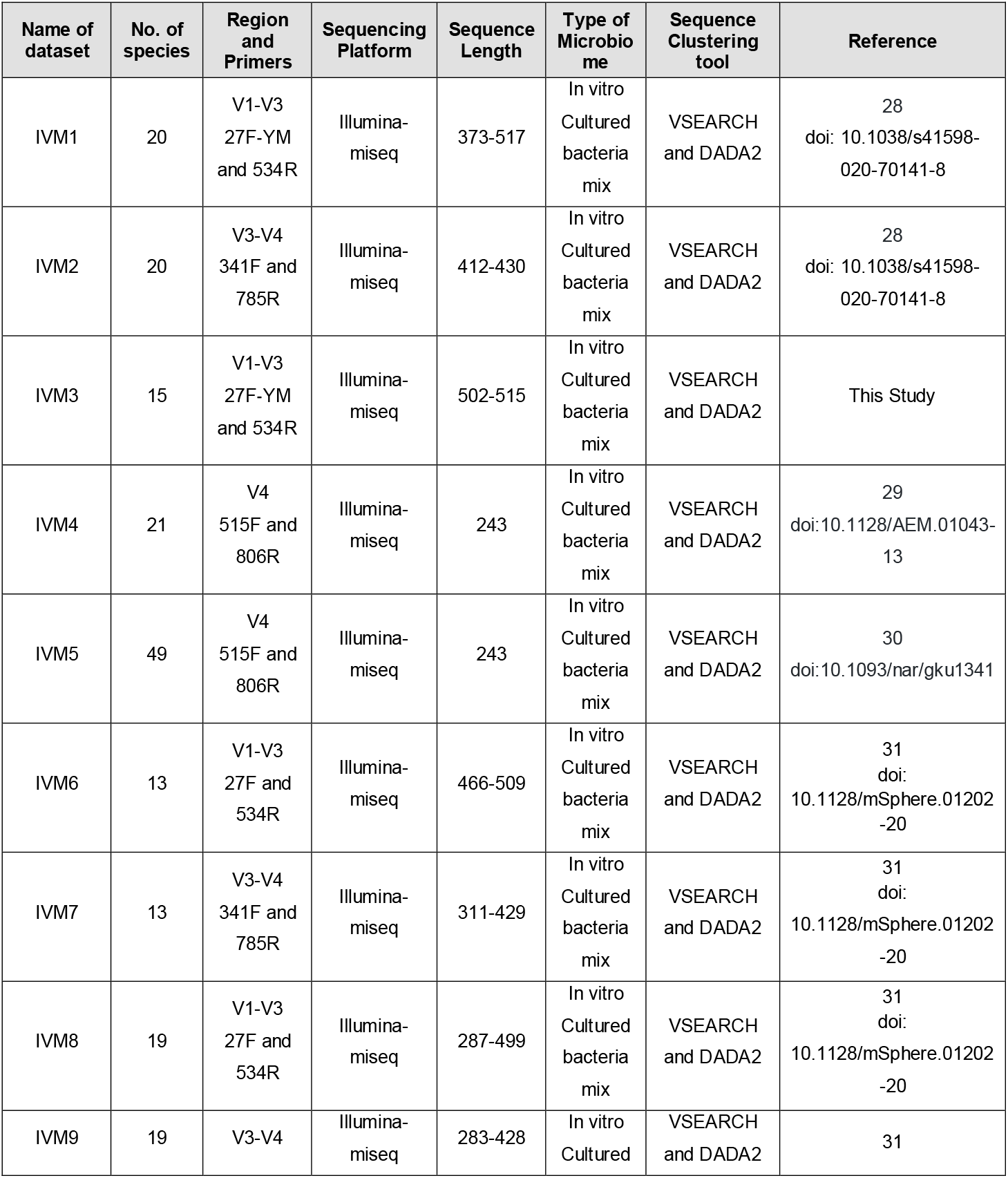
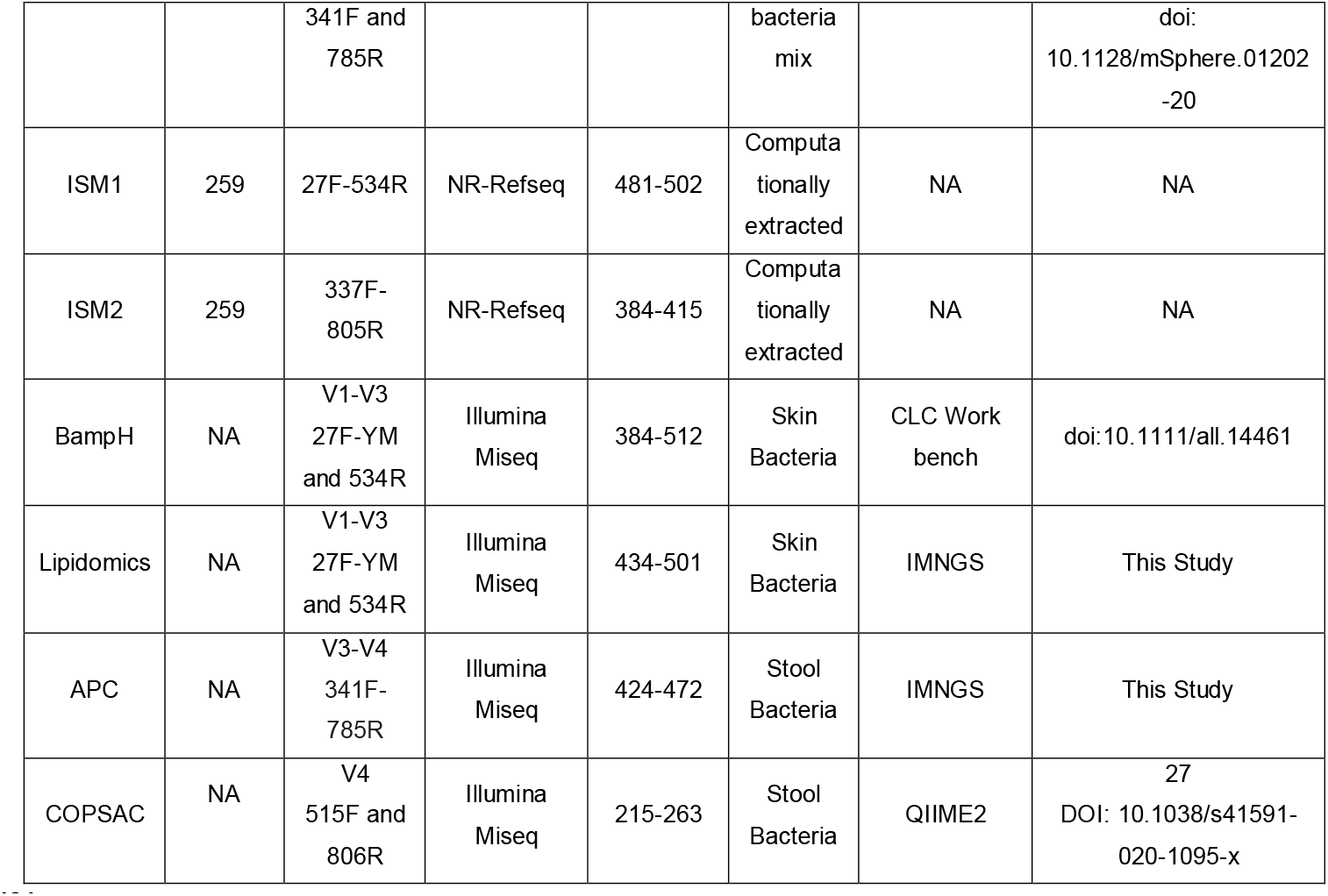
Sequence datasets used for benchmarking.

### Supplementary Figure Captions

**Figure S1:**
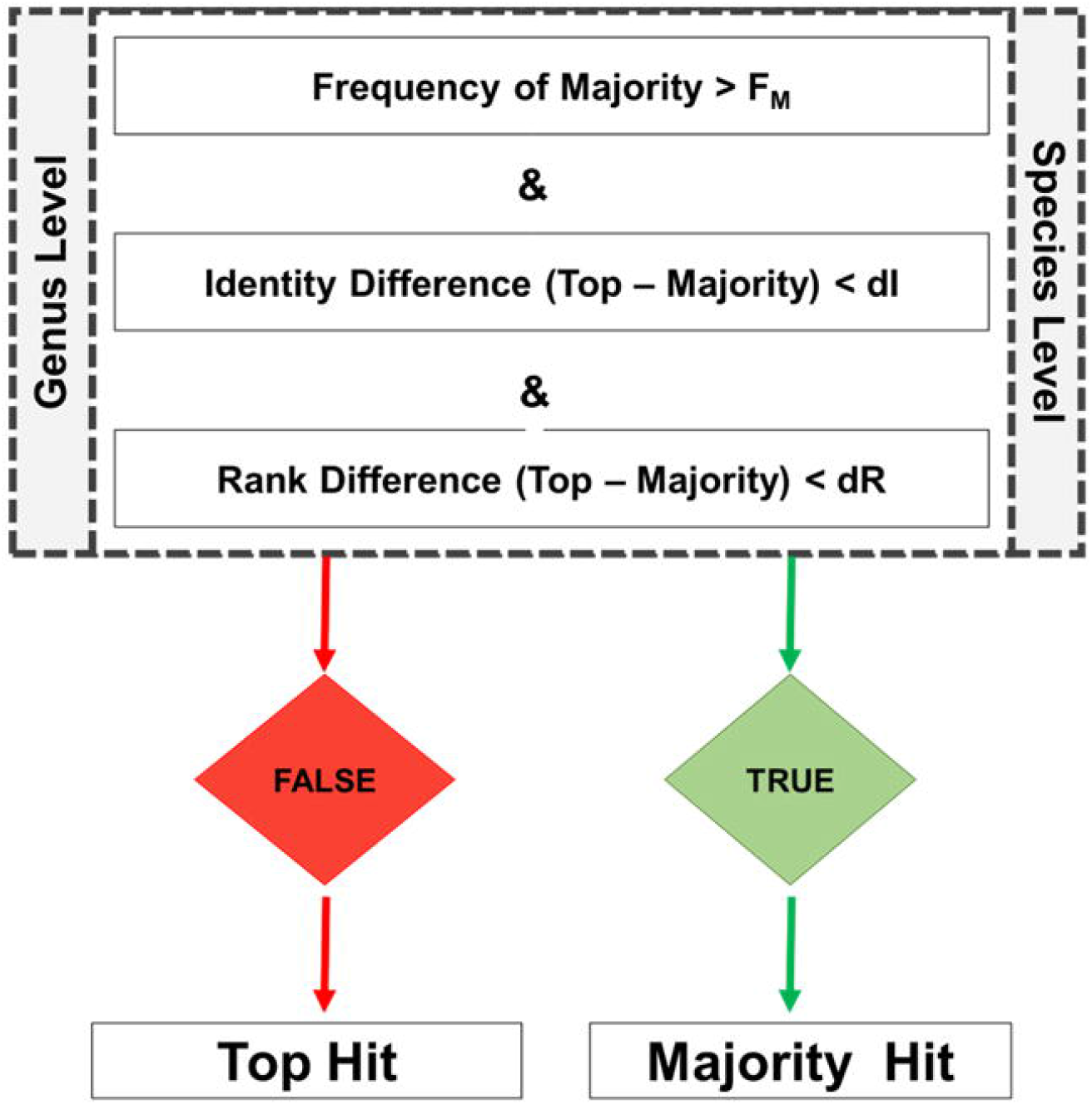
Algorithm used by the AnnotIEM tool for selection of final annotation. Either for species-level annotation, if available, or for genus-level annotation, otherwise, the selection of top-hit versus majority-hit is based on testing 3 parameters against given thresholds (see methods). If the frequency of the same majority-hit species (or genus) is larger than F_M_, and the difference of identity between top-hit and majority-hit is smaller than dI, and the difference of hit rank between top-hit and majority-hit is smaller than dR, then the majority-hit is selected, otherwise the top-hit is selected.

**Figure S2:**
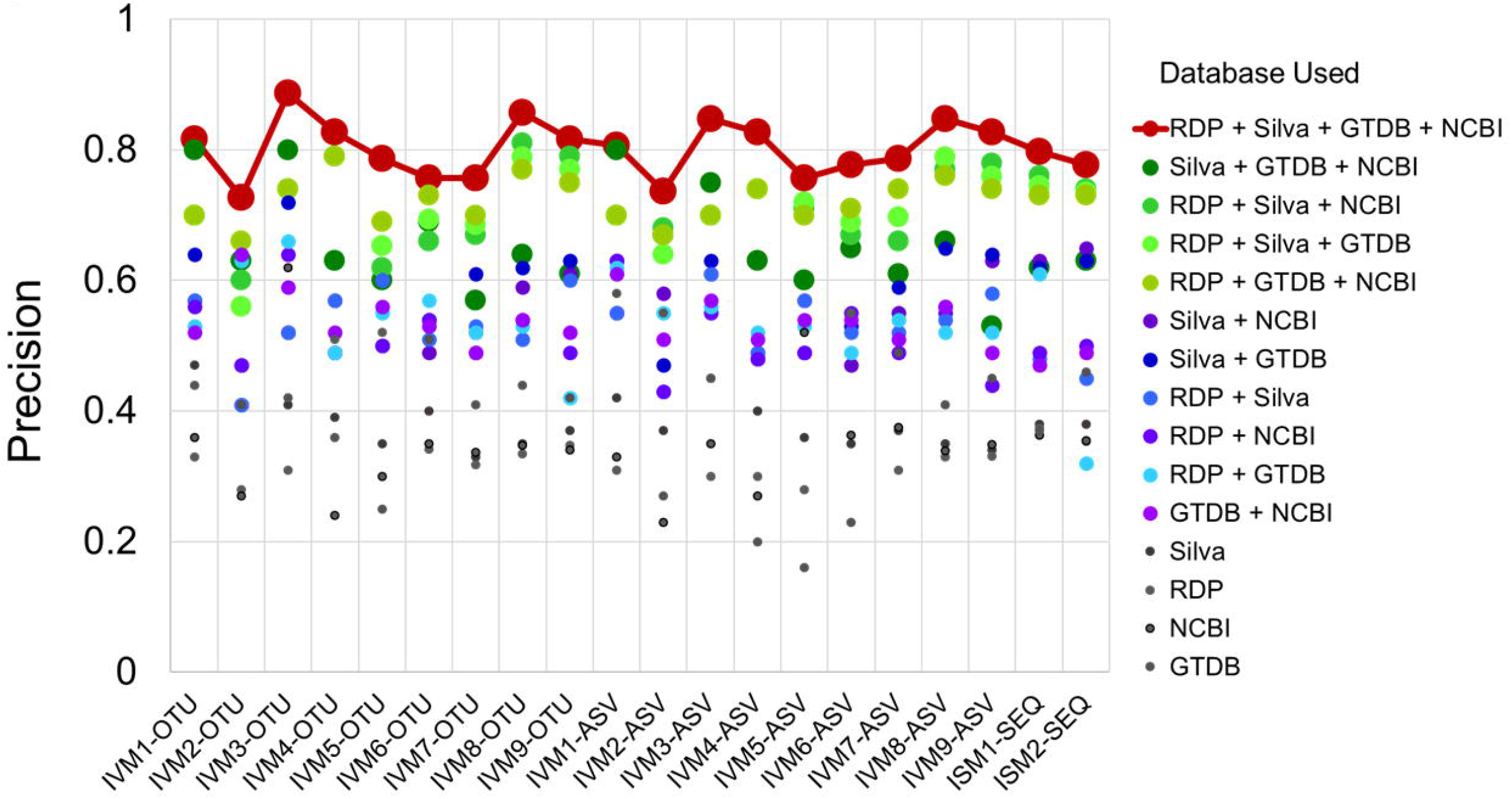
Precision of species-level annotation by AnnotIEM as function of the number of databases used. Precision of the AnnotIEM annotation on species-level in 9 in-vitro mock datasets (IVM), tested both as ASVs and OTUs, and 2 in-silico mock (ISM) datasets (Table 2 and Supplementary Table S1), as function of specific database used or the combination of a number (2, 3 or 4) databases used together. Values achieved by the use of four databases were connected by a line for visualization.

**Figure S3:**
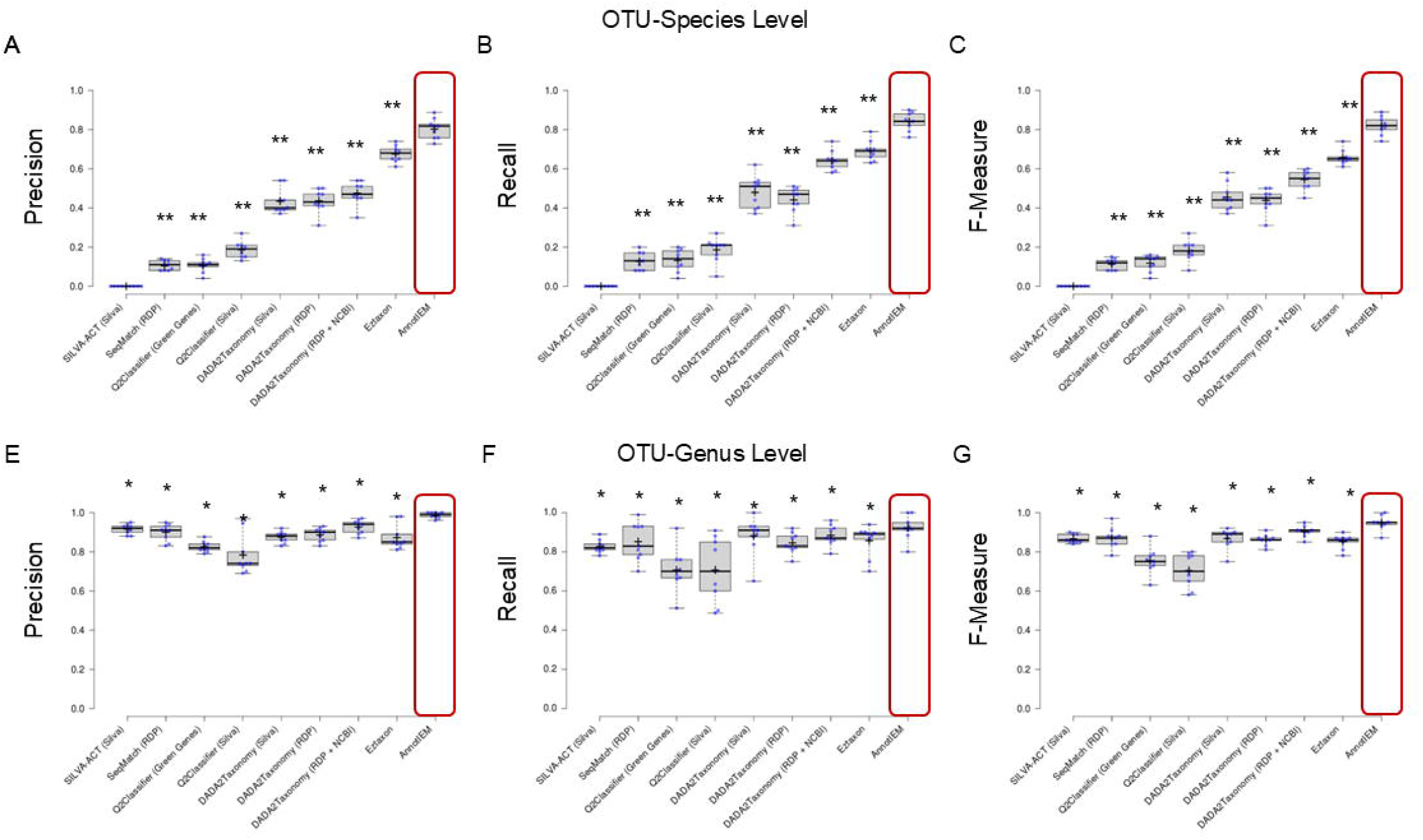
Benchmarking the annotation precision of AnnotIEM against other commonly used annotation tools using OTUs from mock communities. The performance of AnnotIEM is benchmarked against the most widely used annotation tools, using OTUs derived from 9 in-vitro mock (IVM) datasets (Table 2 and Supplementary Table S1). Scoring of the annotation on species-level (A-C) and on genus-level (D-F) is done by Precision (A and D, accordingly), Recall (B and E, accordingly) and F-Measure (C and F, accordingly) scores. The distribution of scores is shown as a box-whisker and scatter plot for each tool. Each box represents the 25th to 75th percentile of the distribution and whiskers extend to the 5th to 95th percentile. The mean is marked by a cross sign, and the median is marked by a horizontal line. * depicts p-value of p<0.05. ** depicts p-value of p<0.01. Commonly used tools used for benchmarking were: SILVA-ACT, SeqMatch RDP, Q2Classifier from QIIME2 using either the Greengenes or the SILVA database, DADA2taxonomy using either SILVA, RDP, or RDP+NCBI databases, and EzBioCloud.

**Figure S4:**
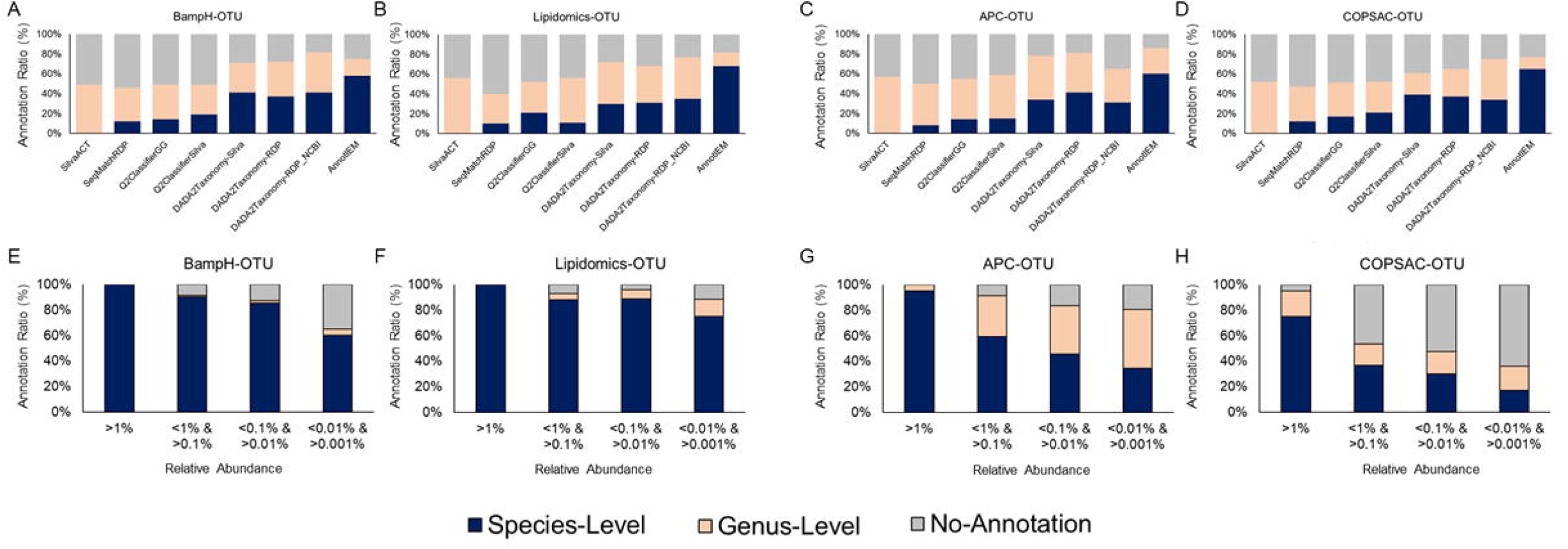
Benchmarking the species-level annotation coverage of AnnotIEM against other commonly used annotation tools using OTUs from real-world microbiome studies. The species-level, and genus-level, annotation coverage (fraction of sequences with species-level, and genus-level, annotation) of AnnotIEM is benchmarked against the most widely used annotation tools, using OTUs derived from 4 real-world large skin or gut microbiome case studies: BampH (A), Lipidomics (B), APC (C), and COPSAC (D), see additional information in Table 2 and Supplementary Table S1. For each of the studies, the species-level and genus-level annotation coverage by AnnotIEM is also depicted as function of the OTUs relative abundance (E-H). Commonly used tools used for benchmarking were: SILVA-ACT, SeqMatch RDP, Q2Classifier from QIIME2 using either the Greengenes or the SILVA database, and DADA2taxonomy using either SILVA, RDP or RDP+NCBI databases. EzBioCloud tool was not benchmarked here because of its limitation of sequencing only one sequence at a time in its free version.

## References

1 Galloway-Peña, J. & Hanson, B. Tools for Analysis of the Microbiome. Dig Dis Sci 65, 674–685, doi:10.1007/s10620-020-06091-y (2020).

2 Maidak, B. L. et al. The RDP (Ribosomal Database Project). Nucleic Acids Res 25, 109–111, doi:10.1093/nar/25.1.109 (1997).

3 Quast, C. et al. The SILVA ribosomal RNA gene database project: improved data processing and web-based tools. Nucleic Acids Res 41, D590–596, doi:10.1093/nar/gks1219 (2013).

4 DeSantis, T. Z. et al. Greengenes, a chimera-checked 16S rRNA gene database and workbench compatible with ARB. Appl Environ Microbiol 72, 5069–5072, doi:10.1128/aem.03006-05 (2006).

5 Yoon, S. H. et al. Introducing EzBioCloud: a taxonomically united database of 16S rRNA gene sequences and whole-genome assemblies. Int J Syst Evol Microbiol 67, 1613–1617, doi:10.1099/ijsem.0.001755 (2017).

6 Parks, D. H. et al. GTDB: an ongoing census of bacterial and archaeal diversity through a phylogenetically consistent, rank normalized and complete genome-based taxonomy. Nucleic Acids Res 50, D785–d794, doi:10.1093/nar/gkab776 (2022).

7 O’Leary, N. A. et al. Reference sequence (RefSeq) database at NCBI: current status, taxonomic expansion, and functional annotation. Nucleic Acids Res 44, D733–745, doi:10.1093/nar/gkv1189 (2016).

8 Edgar, R. C. Search and clustering orders of magnitude faster than BLAST. Bioinformatics 26, 2460–2461, doi:10.1093/bioinformatics/btq461 (2010).

9 Callahan, B. J. et al. DADA2: High-resolution sample inference from Illumina amplicon data. Nat Methods 13, 581–583, doi:10.1038/nmeth.3869 (2016).

10 Landemaine, L. et al. Staphylococcus epidermidis isolates from atopic or healthy skin have opposite effect on skin cells: potential implication of the AHR pathway modulation. Front Immunol 14, 1098160, doi:10.3389/fimmu.2023.1098160 (2023).

11 Severn, M. M. et al. The Ubiquitous Human Skin Commensal Staphylococcus hominis Protects against Opportunistic Pathogens. mBio 13, e0093022, doi:10.1128/mbio.00930-22 (2022).

12 De Tomassi, A. et al. Combining 16S Sequencing and qPCR Quantification Reveals Staphylococcus aureus Driven Bacterial Overgrowth in the Skin of Severe Atopic Dermatitis Patients. Biomolecules 13, doi:10.3390/biom13071030 (2023).

13 Piovani, D. et al. Environmental Risk Factors for Inflammatory Bowel Diseases: An Umbrella Review of Meta-analyses. Gastroenterology 157, 647-659.e644, doi:10.1053/j.gastro.2019.04.016 (2019).

14 Nomura, R. et al. Contribution of Severe Dental Caries Induced by Streptococcus mutans to the Pathogenicity of Infective Endocarditis. Infect Immun 88, doi:10.1128/iai.00897-19 (2020).

15 Tong, H., Gao, X. & Dong, X. Streptococcus oligofermentans sp. nov., a novel oral isolate from caries-free humans. Int J Syst Evol Microbiol 53, 1101–1104, doi:10.1099/ijs.0.02493-0 (2003).

16 Bolyen, E. et al. Reproducible, interactive, scalable and extensible microbiome data science using QIIME 2. Nat Biotechnol 37, 852–857, doi:10.1038/s41587-019-0209-9 (2019).

17 Lagkouvardos, I. et al. IMNGS: A comprehensive open resource of processed 16S rRNA microbial profiles for ecology and diversity studies. Sci Rep 6, 33721, doi:10.1038/srep33721 (2016).

18 Schloss, P. D. et al. Introducing mothur: open-source, platform-independent, community-supported software for describing and comparing microbial communities. Appl Environ Microbiol 75, 7537–7541, doi:10.1128/aem.01541-09 (2009).

19 Bokulich, N. A. et al. Optimizing taxonomic classification of marker-gene amplicon sequences with QIIME 2’s q2-feature-classifier plugin. Microbiome 6, 90, doi:10.1186/s40168-018-0470-z (2018).

20 Pruesse, E., Peplies, J. & Glöckner, F. O. SINA: accurate high-throughput multiple sequence alignment of ribosomal RNA genes. Bioinformatics 28, 1823–1829, doi:10.1093/bioinformatics/bts252 (2012).

21 Cole, J. R. et al. Ribosomal Database Project: data and tools for high throughput rRNA analysis. Nucleic Acids Res 42, D633–642, doi:10.1093/nar/gkt1244 (2014).

22 Camacho, C. et al. BLAST+: architecture and applications. BMC Bioinformatics 10, 421, doi:10.1186/1471-2105-10-421 (2009).

23 Suzuki, H., Lefébure, T., Bitar, P. P. & Stanhope, M. J. Comparative genomic analysis of the genus Staphylococcus including Staphylococcus aureus and its newly described sister species Staphylococcus simiae. BMC Genomics 13, 38, doi:10.1186/1471-2164-13-38 (2012).

24 Johnson, J. S. et al. Evaluation of 16S rRNA gene sequencing for species and strain-level microbiome analysis. Nat Commun 10, 5029, doi:10.1038/s41467-019-13036-1 (2019).

25 Olendraite, I., Brown, K. & Firth, A. E. Identification of RNA Virus-Derived RdRp Sequences in Publicly Available Transcriptomic Data Sets. Mol Biol Evol 40, doi:10.1093/molbev/msad060 (2023).

26 Abellan-Schneyder, I. et al. Primer, Pipelines, Parameters: Issues in 16S rRNA Gene Sequencing. mSphere 6, doi:10.1128/mSphere.01202-20 (2021).

27 Kozich, J. J., Westcott, S. L., Baxter, N. T., Highlander, S. K. & Schloss, P. D. Development of a dual-index sequencing strategy and curation pipeline for analyzing amplicon sequence data on the MiSeq Illumina sequencing platform. Appl Environ Microbiol 79, 5112–5120, doi:10.1128/aem.01043-13 (2013).

28 Schirmer, M. et al. Insight into biases and sequencing errors for amplicon sequencing with the Illumina MiSeq platform. Nucleic Acids Res 43, e37, doi:10.1093/nar/gku1341 (2015).

29 Soriano-Lerma, A. et al. Influence of 16S rRNA target region on the outcome of microbiome studies in soil and saliva samples. Sci Rep 10, 13637, doi:10.1038/s41598-020-70141-8 (2020).

30 Stokholm, J. et al. Maturation of the gut microbiome and risk of asthma in childhood. Nat Commun 9, 141, doi:10.1038/s41467-017-02573-2 (2018).

31 Hülpüsch, C. et al. Skin pH-dependent Staphylococcus aureus abundance as predictor for increasing atopic dermatitis severity. Allergy 75, 2888–2898, doi:10.1111/all.14461 (2020).

