## Supplemental Table 1 for "AnnotIEM: novel tool for microbiome species-level annotation of 16S gene based microbial sequencing"

**Supplementary Table S1: Sequence datasets used for benchmarking.**

| Name of dataset | No. of species | Region and Primers | Sequencing Platform | Sequence Length | Type of Microbiome | Sequence Clustering tool | Reference |
|---|---|---|---|---|---|---|---|
| IVM1 | 20 | V1-V3<br>27F-YM and 534R | Illumina-miseq | 373-517 | In vitro Cultured bacteria mix | VSEARCH and DADA2 | 28<br>doi: 10.1038/s41598-020-70141-8 |
| IVM2 | 20 | V3-V4<br>341F and 785R | Illumina-miseq | 412-430 | In vitro Cultured bacteria mix | VSEARCH and DADA2 | 28<br>doi: 10.1038/s41598-020-70141-8 |
| IVM3 | 15 | V1-V3<br>27F-YM and 534R | Illumina-miseq | 502-515 | In vitro Cultured bacteria mix | VSEARCH and DADA2 | This Study |
| IVM4 | 21 | V4<br>515F and 806R | Illumina-miseq | 243 | In vitro Cultured bacteria mix | VSEARCH and DADA2 | 29<br>doi:10.1128/AEM.01043-13 |
| IVM5 | 49 | V4<br>515F and 806R | Illumina-miseq | 243 | In vitro Cultured bacteria mix | VSEARCH and DADA2 | 30<br>doi:10.1093/nar/gku1341 |
| IVM6 | 13 | V1-V3<br>27F and 534R | Illumina-miseq | 466-509 | In vitro Cultured bacteria mix | VSEARCH and DADA2 | 31<br>doi: 10.1128/mSphere.01202-20 |
| IVM7 | 13 | V3-V4<br>341F and 785R | Illumina-miseq | 311-429 | In vitro Cultured bacteria mix | VSEARCH and DADA2 | 31<br>doi: 10.1128/mSphere.01202-20 |
| IVM8 | 19 | V1-V3<br>27F and 534R | Illumina-miseq | 287-499 | In vitro Cultured bacteria mix | VSEARCH and DADA2 | 31<br>doi: 10.1128/mSphere.01202-20 |
| IVM9 | 19 | V3-V4<br>341F and 785R | Illumina-miseq | 283-428 | In vitro Cultured bacteria mix | VSEARCH and DADA2 | 31<br>doi: 10.1128/mSphere.01202-20 |
| ISM1 | 260 | 27F-534R | NR-Refseq | 481-502 | Computationally extracted | NA | NA |

| | | | | | | | |
|---|---|---|---|---|---|---|---|
| ISM2 | 260 | 337F-805R | NR-Refseq | 384-415 | Computationally<br>extracted | NA | NA |
| BampH | NA | V1-V3<br>27F-YM and 534R | Illumina Miseq | 384-512 | Skin Bacteria | CLC Work<br>bench | doi:10.1111/all.14461 |
| Lipidomics | NA | V1-V3<br>27F-YM and 534R | Illumina Miseq | 434-501 | Skin Bacteria | IMNGS | This Study |
| APC | NA | V3-V4<br>341F-785R | Illumina Miseq | 424-472 | Stool Bacteria | IMNGS | This Study |
| COPSAC | NA | V4<br>515F and 806R | Illumina Miseq | 215-263 | Stool Bacteria | QIIME2 | 27<br>DOI: 10.1038/s41591-020-1095-x |

**Formatted:** Centered
